# Point mutations of the mitochondrial chaperone TRAP1 affect its functions and pro-neoplastic activity

**DOI:** 10.1101/2024.10.24.619840

**Authors:** Claudio Laquatra, Alessia Magro, Federica Guarra, Matteo Lambrughi, Giulio Fracasso, Melissa Bacchin, Lavinia Ferrone, Martina La Spina, Elisabetta Moroni, Elena Papaleo, Giorgio Colombo, Andrea Rasola

## Abstract

The mitochondrial chaperone TRAP1 is a key regulator of cellular homeostasis and its activity has important implications in neurodegeneration, ischemia and cancer. Recent evidence has indicated that TRAP1 mutations are involved in several disorders, even though the structural basis for the impact of point mutations on TRAP1 functions has never been studied. By exploiting a modular structure-based framework and molecular dynamics simulations, we investigated the effect of five TRAP1 mutations on its structure and stability. Each mutation differentially impacts long-range interactions, intra and inter-protomer dynamics and ATPase activity. Changes in these parameters influence TRAP1 functions, as revealed by their effects on the activity of the TRAP1 interactor succinate dehydrogenase (SDH). In keeping with this, TRAP1 point mutations affect the growth and migration of aggressive sarcoma cells, and alter sensitivity to a selective TRAP1 inhibitor. Our work provides new insights on the structure-activity relationship of TRAP1, identifying crucial amino acid residues that regulate TRAP1 proteostatic functions and pro-neoplastic activity.

## Introduction

The molecular chaperone TRAP1 is a mitochondrial member of the heat shock protein 90 (HSP90) family that assists the folding of a variety of interacting proteins called clients, tuning their stability and activity (1, 2). The TRAP1 chaperone cycle is coupled with ATP binding and hydrolysis in the N-terminal domain (NTD), whereas the middle domain (MD) controls ATP hydrolysis and client binding and the C-terminal domain (CTD) mediates the dimerization of the two protomers required for TRAP1 activity(3, 4). ATP binding induces buckling in one of the two protomers, and the subsequent hydrolysis transfers this asymmetry to the second monomer, activating its ATPase function and completing the cycle(5). This two-step process facilitates proper client remodeling and release, with the dimers flipping from a closed to an open (*apo*) conformation, in which they reset for a new chaperone cycle(5–7). Hence, the conformational changes elicited by ATP binding and hydrolysis are intimately linked to TRAP1 internal dynamics, regulating its interactions with clients and shaping its activity (4, 8). In this scenario, the fluctuations of residues on short timescales have been shown to facilitate slow and large-scale motions that are linked to chaperone functions. However, it remains unclear how TRAP1 ATPase and chaperone cycle finely tune client interaction and activity and determine specific biochemical and biological outputs.

In tumors, TRAP1 is involved in the protection from oxidative stress(9, 10), cell death inhibition by preventing mitochondrial permeability transition(11, 12) and induction of a metabolic switch toward aerobic glycolysis by down-regulating the activity of the OXPHOS components cytochrome *c* oxidase and succinate dehydrogenase (SDH), thus eliciting succinate-dependent stabilization of the hypoxia-inducible factor HIF1α(13–16). As a result, TRAP1 boosts an aggressive pseudo-hypoxic phenotype, and its increased expression correlates with disease progression and poor prognosis in diverse cancer types(4), whereas TRAP1 genetic or pharmacological inhibition impedes the growth of various neoplastic models(16–18). It has been shown that post-translational modifications like S-nitrosylation, phosphorylation, and acetylation/deacetylation can modulate TRAP1 activity, influencing its eventual effects on tumor growth(13, 16, 19–22). Moreover, TRAP1 point mutations have been identified in several pathological conditions including Parkinson’s disease, autoinflammatory diseases associated with redox disequilibrium, congenital anomalies of the kidney and urinary tract (CAKUT), and children with the mitochondrial disease Leigh Syndrome and with a group of growth abnormalities called VECTERL syndrome(23–26). Taken together, these findings suggest that alterations in the structure of TRAP1 may lead to changes in its functions and activity. However, a thorough examination of the structure-activity relationship of TRAP1 mutations and their potential effects on its biological functions and on disease progression has not yet been conducted, particularly in the context of tumors. Here, by applying a step-wise approach encompassing bioinformatic predictions*, in vitro* and *in cellulo* studies, we have identified a set of point mutations that confer unique properties to the dynamics, stability, activity and pro-tumorigenic functions of TRAP1.

## Results

### High throughput screening and classification of TRAP1 point mutations

In order to reveal TRAP1 point mutations and to provide indications on their effect on its activity, we first performed a bulk-tissue gene-expression analysis that revealed widespread but heterogeneous TRAP1 expression, both in normal tissues and in a variety of cancer types (**Supplementary Fig. 1a-b**). We then identified 310 TRAP1 variants and utilized the MAVISp (Multi-layered Assessment of Variants by Structure for Proteins) structure-based computational framework (27) to characterize their effects, classifying them based on predicted stability and pathogenicity (**Fig. 1a, Supplementary fig. 1c and additional file 1**). MAVISp analyzes variants based on 3D structures, interaction networks, and evolutionary data to predict the impact of mutations on protein stability and pathogenicity, particularly in relation to diseases (27). We then selected five variants according to their frequency, pathogenicity score (p.s.) and location within the TRAP1 structure (**Fig. 1b-c**). In detail, the variant D260N is located in the TRAP1 NTD that harbors the Mg^2+^ and ATP/ADP binding sites; three other variants (P381S, V556M, A571T) are in the MD, which contains the binding site for client proteins, while T600P is in the CTD that regulates TRAP1 dimerization (**Fig. 1c**). For each of these mutations, we extracted data from the MAVISp database regarding their predicted effects on structural stability, phosphorylation status and long-range impacts. Additionally, we assessed the effects of these variants on TRAP1 homodimerization and their local and distal effects on TRAP1 client binding sites (**Fig. 1d**). Three mutations (P381S, V556M, T600P) showed damaging effects in MAVISp to some extent; P381S displayed the highest pathogenicity score (p.s. 808) and T600P a destabilizing effect on TRAP1 structure (**Fig. 1d**).

**Figure 1.**
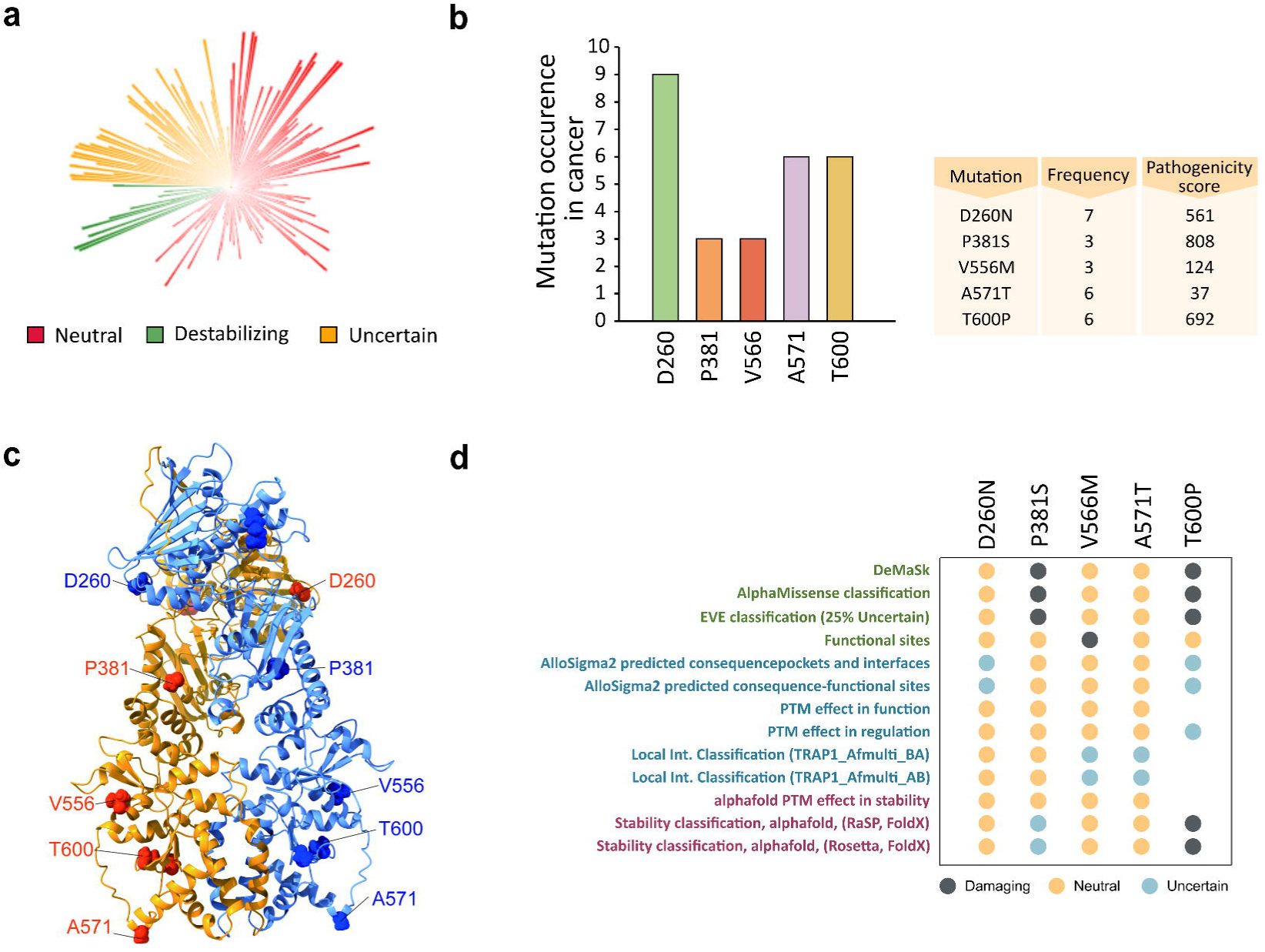
High throughput screening and classification of TRAP1 point mutations. **a** Circular blot indicating the 310 variants identified for TRAP1 with the MAVISp computational framework. Mutations were stratified according to their effect on protein stability in neutral, uncertain and destabilizing. The height of each histogram indicates the pathogenicity score associated to each variants. **b** Most frequent TRAP1 mutations from COSMIC database and relative table indicating the frequency and pathogenicity score of selected variants. **c** Cartoon representation of the structure of TRAP1 homodimer TRAP170-704 (Alphafold-Multimer model) with spheres highlighting the Cα atoms of the five residues affected by the selected variants. **d** The dot plot illustrates the pathogenicity scores (DeMaSk, AlphaMissense, and EVE) and the predicted mechanisms of the effects of selected variants included in the MAVISp framework.

### Effect of point mutations on TRAP1 stability

Point mutations can significantly affect protein stability, potentially causing misfolding and altering chaperone activity. To investigate this, we evaluated the impact of the selected point mutations on TRAP1 stability. The MAVISp screening identified the T600P substitution as the only mutation with a destabilizing effect. Indeed, T600 is localized at the C-terminal domain and highly buried from the solvent (solvent accessible surface area SASA 6%). The substitution of threonine (T) with proline (P) strongly affects the interaction with the surrounding residues including H558, K598-V599, L601-L603, M609, and E643-P646 (**Fig. 2a**), as well as the hydrogen bond network. To further investigate the effect of the mutation on local interactions, we ran 900 ns long MD simulations (three 300 ns independent replicates) for both wild-type and mutant forms of TRAP1 (see below). For each system, 600 ns made up of the equilibrated parts of the trajectories were analyzed with respect to the WT protein. The MD simulations were based on the structure of wild-type Zebrafish TRAP1 (zTRAP1), whose residue numbering is offset by 15 residues relative to the human protein (see material and methods and **Supplementary Fig. 2a**). In our model, the mutated Thr in protomer A (T615 in zTrap1-WT) is part of a β-sheet for about 51% of the analysed MD trajectory, which is likely to be disrupted by mutation to Pro. Conversely, in protomer B the same β-strand is consistently shorter and the assigned secondary structure for T615 is that of a bend (94% frequency). Overall, Pro alters the H-bond interactions observed for T615 during MD simulations in both protomers in zTrap1 (**Fig. 2b**) and only backbone interactions with the N+2 residue R617 are observed with low occupancy. Occupancy times of the H-bond interactions involving the mutated position are shown for TRAP1 WT and T600P variant in **Supplementary Fig 2b**.

**Figure 2.**
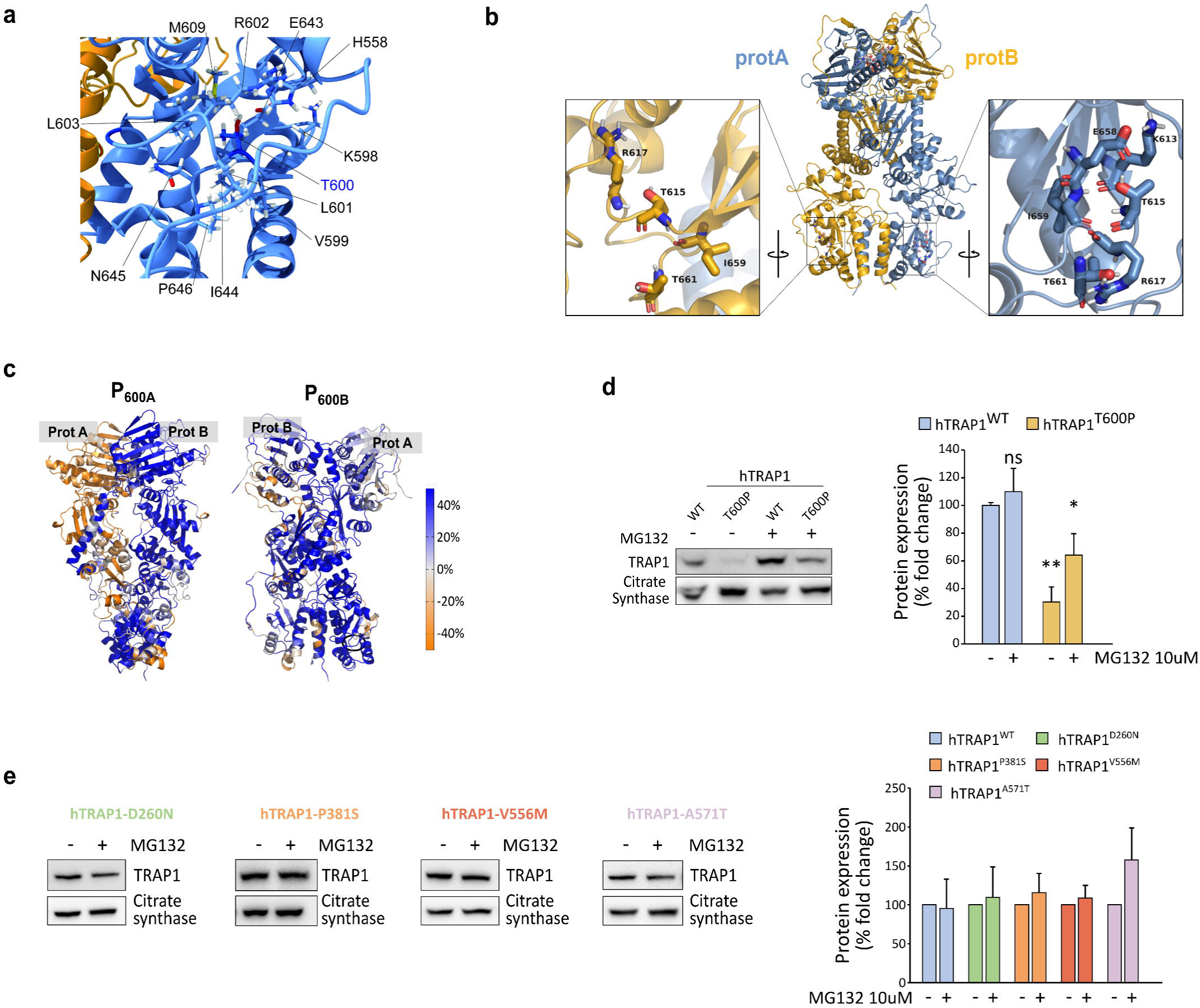
Effect of point mutations on TRAP1 stability. **a** T600 and its surrounding residues are visualized on the 3D structure of the TRAP1 homodimer. **b** Residues surrounding Thr600 in both protomer A (light blue) and B (dark yellow) of TRAP1. In the zoomed views the structure is rotated and centered on the mutated position. Numbering is relative to zTrap1 sequence as in PDB code 4IPE. To convert to hTrap1 numbering 15 should be subtracted. **c** change in DF score to mutated positions in prot. A or B for each residue in going from WT to T600P (indicated by black circles) projected onto the protein 3D structure (P_600A_, P_600B_). Colour code for DF scores: blue areas (positive values) correspond to lower mechanical coordination in Trap1 mutants to the respect of the WT protein, whereas orange ones (negative values) indicate a higher coordination in the first. Grey/white areas are those unaffected by the mutation. **d** Western blot of sMPNST cells expressing the human WT or T600P TRAP1 forms. Where indicated, cells were treated with the proteasomal inhibitor MG132 10 μM for 6 hours. Data are reported as average ± S.E.M. of 3 independent experiments with a two-tail unpaired *t* test. **e** Western blot and relative quantification of sMPNST expressing the indicated mutant forms of TRAP1 treated or not with 10 uM of MG132 for 6 hours. Data are reported as average ± S.E.M. of 3 independent experiments.

Next, starting from MD trajectories, we conducted a pair-wise residue fluctuation analysis (Distance Fluctuations (DF)) to check the impact of the mutated residue on mechanical coordination. We then obtained the percentage difference matrix by subtracting DF_WT_ from DF_T600P_ (%ΔDF, see Methods section and **Supplementary Fig 2c**). By projecting the columns corresponding to the mutated positions in protomers A and B in the 3D protein structure (P_600A_ and P_600B_), we observed the changes in the DF score (mechanical coordination) to the mutated positions of all the other residues from WT to T600P. Residues that lose mechanical coordination (allosteric dialogue) upon mutation are shown in blue, while those with increased coordination are in orange. The predominance of blue indicates a widespread long-range loss of mechanical coordination from WT to T600P (**Fig 2c**). These findings suggest that the destabilization around this residue might affect the entire protein, potentially leading to incorrect unfolding and/or dimer destabilization further supporting the MAVISp prediction. Accordingly, only a faint re-expression of the human T600P TRAP1 (hTRAP1-T600P) could be obtained in murine sarcoma cells where the endogenous TRAP1 had been knocked out(17), while a significant increase in protein expression occurred upon proteasome inhibition (**Fig. 2d**). Conversely, the other mutations did not affect the stability or re-expression of the human TRAP1 variants, as predicted by MAVISp (**Fig. 2e** and **supplementary Fig. 2d**).

### Point mutations have differential effects on TRAP1 ATPase activity, protein dynamics and internal mechanical connectivity

TRAP1 functions depend on conformational changes driven by the protein’s internal dynamics. Substituting specific residues can disrupt local interactions, causing structural deformations over time that affect protein activity. Consistently, previous studies have shown that post-translational modifications or mutations at specific residues of TRAP1 influence its ATPase activity (22), indicating that alterations in TRAP1 structure and dynamics may impact its functions. To verify the predictive effect of point mutations, we evaluated each mutant by measuring i) TRAP1 ATPase activity on purified recombinant proteins and ii) its structural positioning, effects on protein conformations, local interaction networks, and dynamic coordination patterns via pair-residue fluctuation analysis. These combined analyses aim to offer both a quantitative and qualitative understanding of how a single point mutation can disrupt TRAP1’s local and global dynamic properties.

We observed that the hTRAP1-D260N variant shows an increased TRAP1 activity of about 50% with respect to the WT protein (**Fig. 3a**) The residue D260 is in a solvent-exposed region of the NTD, behind the ATP active site and close to the dimer interface where the NTD of one protomer contacts the MD of the other. The substitution of D to N disrupts native H-bond interactions. Indeed, D260 (D275 in zTrap1) is closely positioned to K124 and K273 in both protomers, and D260 of protomer B is near the R427 of protomer A. In the D260N mutant, these interactions are significantly reduced or absent (**Supplementary Fig. 3a**). The K124 is located at the N terminus of the α-helix 2 (residues 123-149) (28), which includes the ATP binding site and residues crucial for ATP binding and hydrolysis (the catalytic E130 and Y121, R129, S133) (29). Instead, R427 (prot. B) is in the α-helix 13 (residues 423-448), close to the NTD-MD dimer interface and adjacent to R417, which directly binds ATPγ phosphate, acting as an ATP sensor and stabilizing the closed conformation required for ATP hydrolysis (6, 28, 30). The mutation D260N also affects protein dynamics as an increase in the overall mechanical coordination is observed in mutated TRAP1 with respect to the WT protein (P_260A/B_, **Fig. 3b and additional file 2**). Accordingly, the analysis of the number of mechanically connected residues along the sequence (mechanical connectivity index η, see Methods section) and its variation relative to the WT TRAP1 (Δη_D260N_) reveals a general increase for both protomer NTDs (**Fig. 3c and Supplementary Fig 3c**). Moreover, the increased mechanical connectivity P_260A/B_ and the mechanical connectivity index of NTDs suggest greater rigidity and efficient allosteric communication in ATP hydrolysis areas, consistent with a higher ATPase rate observed for this mutant.

**Figure 3.**
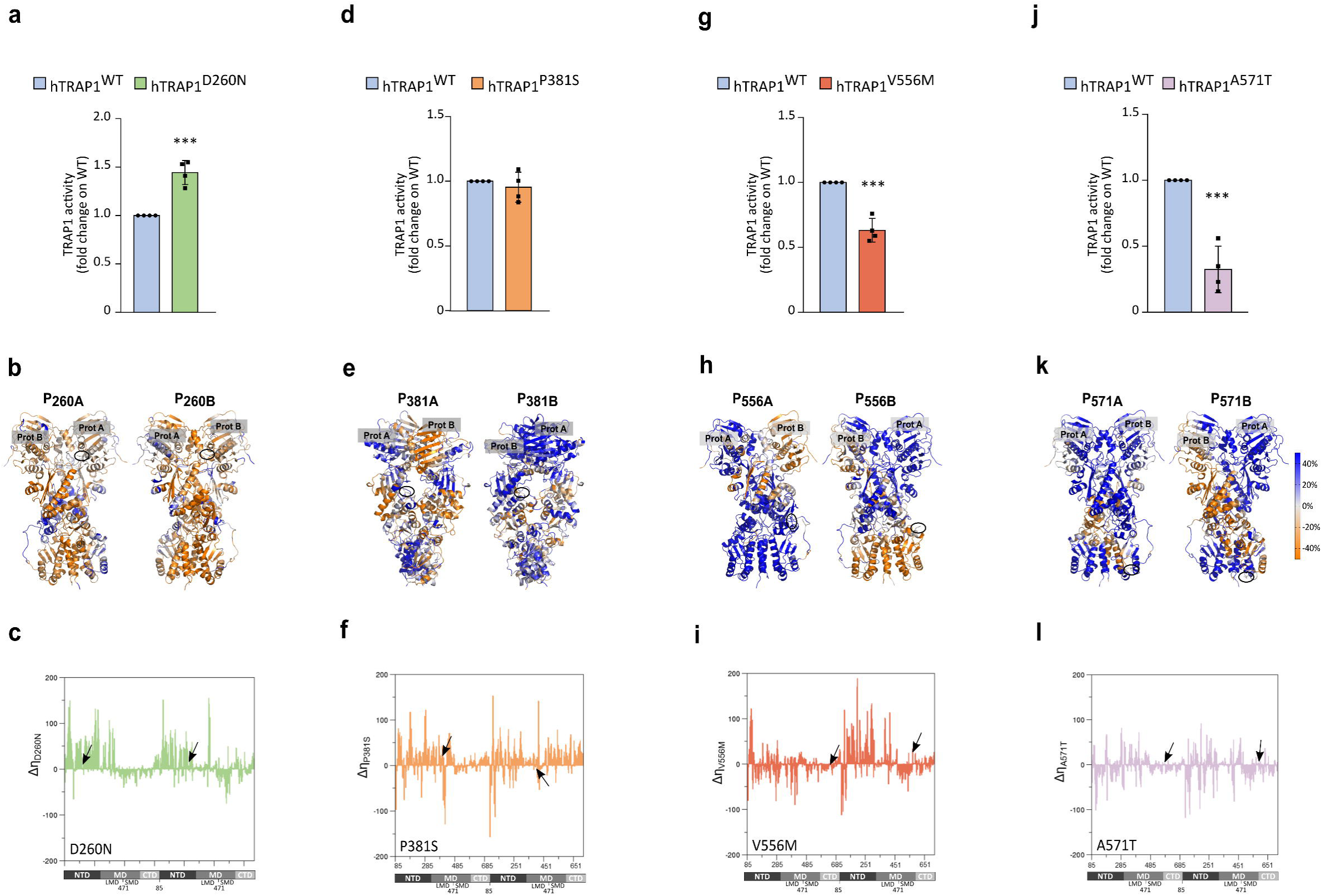
Effect of point mutations on TRAP1 ATPase activity and molecular dynamics. **a**, **d, g** and **j** ATPase activity of recombinant WT or mutant forms of human TRAP1 was measured as PO_4_^3-^ released. Data are shown as fold change (with respect to the WT protein) and represent the mean ± SEM of n=4 independent experiments done in triplicate. **b, e, h** and **k** Change in DF score to mutated positions in prot. A or B for each residue from WT to Mut Trap1 (**b**, D260N; **e** P381S; **h** V556M; **k** A571T), projected onto the protein 3D structure (P_mutA_, P_mutB_). Protein views are rotated to allow a visualization of the mutation position, the identity of the protomer is labelled. Colour code for DF scores: blue areas (positive values) correspond to lower mechanical coordination in Trap1 mutants to the respect of WT protein, whereas orange ones (negative values) indicate a higher coordination in the first. Grey/white areas are those unaffected by the mutation. **c, f, i, l** Variation in the mechanical connectivity index for each mutant along the sequence (Δη_mut_). On the x axis numbering of zTrap1 as in PDB code 4IPE is shown together with the corresponding domains of prot. A and prot.B. and the residue numbering is from 85 to 719. NTD: N-terminal Domain, residues 85-310; MD: Middle Domain divided in the subdomains Large Middle Domain (LMD), residues 311-470 and Small Middle Domain (SMD) residues 471-586; CTD: C-Terminal Domain, residues 587-719.

P381S mutation does not result in a significant change in the activity of the recombinant protein if compared with hWT-TRAP1 (**Fig. 3d**). This residue is located in the Large Middle Domain (LMD) in proximity of the lumen of the protein at the dimer interface, where unfolded portions of clients bind in Hsp90 (PDB code 7KCM and (31)). The mutation from Pro to Ser increases the number of H-bond interactions involving the mutated residue (**Supplementary Fig 3b**) and partially enhances mechanical coordination in protomer A, although this variant causes an overall trend of reduced coordination (P_381A/B_, **Fig. 3e and additional file 2**). The mechanical connectivity index of this variant is similar to D260N but shows a smaller increase in the NTDs. However, the MD and CTD of protomer B have an overall increase in mechanically connected residues, unlike other variants (**Fig. 3f and Supplementary Fig 3c**). Even if the mechanical connectivity index profile is similar to that of the D260N variant, the increase in NTDs is less pronounced and in line with a functional protein with an ATPase activity comparable to the one of WT-TRAP1. Interestingly, an increase in the dynamic coordination in this region can be predicted to favour a more efficient remodelling of client proteins.

The mutant V556M displays a reduction in ATPase activity if compared to wild-type TRAP1 (**Fig. 3g**). This residue is in the C-terminal Domain (CTD) at the border with the Small Middle Domain (SMD) and belongs to a disordered flexible linker in protomer A (not solved in 4IPE crystal structure); while the same region is structured into an α helix in protomer B. Concerning the internal dynamics of the protein, there is a general decrease in the mechanical coordination to the mutated position (P_556A/B_ in **Fig. 3h and additional file 2**). The mechanical connectivity increases in NTD of protomer B but generally decreases in the rest of the protein. This includes the entire buckled protomer A, which undergoes the first ATP hydrolysis and the SMD of protomer B, which includes the mutated position and is involved in client binding (**Fig. 3i and Supplementary Fig 3c**). These observations support the observed reduction in TRAP1 activity.

Finally, we analyzed the variant A571T finding a marked reduction in ATPase activity compared to wild-type TRAP1 (**Fig. 3j**). Ala 571 (Serine in WT zTrap1) is located in the CTD within the loop regions. In protomer A, it is part of an unstructured loop (not modelled in the 4IPE structure), while in protomer B, it is part of a short helix (310 or α) for about 38% of the MD trajectory or turn/bend (frequency 62%). Interestingly, when Ala is mutated to Thr, the frequency of the helix fold in protomer B significantly increases to 78%, while the bend frequency in protomer A rises to 5%, indicating a substantial impact of the mutation on the secondary structure of this region. Therefore, we examined the effect of this mutation on the protein’s overall internal dynamics through pair residue fluctuation analysis (DF analysis). Similar to the Trap1 V556M mutation, we observed a general decrease in mechanical coordination at the mutated position (P_571A/B_ **Fig.3k and additional file 2**). Consistent with this finding, the Δη graph shows an overall flattening compared to the other mutants, with a prevalence of negative peaks. This indicates a decrease in the mechanical connectivity index η from WT to A571T protein, particularly in the CTDs where the mutation is located (**Fig. 3lv and Supplementary Fig. 3c**).

Overall, our results suggest a model in which the introduction of specific mutations disrupts the conformational profile of TRAP1, altering its activity, dynamic states and sensitivity to the selective inhibitor.

### Point mutations affect TRAP1 bioenergetic properties

We previously showed that TRAP1 downregulates SDH activity causing the succinate-dependent stabilization of HIF1α and the consequent neoplastic cell growth (14–16). Therefore, we used mouse Malignant Peripheral Nerve Sheath Tumor cells (sMPNST cells) where TRAP1 plays a crucial pro-neoplastic role to assess how the different TRAP1 variants tune SDH activity. By re-expressing either the human wild-type or mutant forms of TRAP1 upon genetic silencing of the endogenous protein (**Supplementary fig. 2d**), we found the expected increase in the succinate-coenzyme Q reductase (SQR) activity of SDH following ablation of TRAP1 expression, which was rescued to the level of parental cells upon re-expression of the h-TRAP1-WT (**Fig. 4a**). Interestingly, cells expressing either h-TRAP1-P381S or h-TRAP1-D260N display the same SQR activity of cells harboring h-TRAP1-WT (**Fig. 4b**), while re-expression of hTRAP1-V556M and hTRAP1-A571T increases SQR activity to the same extent reached following genetic ablation of TRAP1 (**Fig. 4b**). Notably, expression of TRAP1 variants or absence of the endogenous protein does not affect SDHA protein levels (**Fig. 4c**). The residue V556 is located in the the C-terminal Domain (CTD) close to the Small Middle Domain (SMD). This region in protomer B is shown to interact with the model client protein SdhB in a reported Cryo-EM structure (PDB code 7KCM), with V556 being directly in contact with the client (**Fig. 4d**). Focusing on the dynamics of the region surrounding the mutated position, some of which contact SdhB (residues 521-642, protomer B), we observed an overall increase in the flexibility upon mutation, as demonstrated by RMSF analysis (**Fig. 4e**).

**Figure 4.**
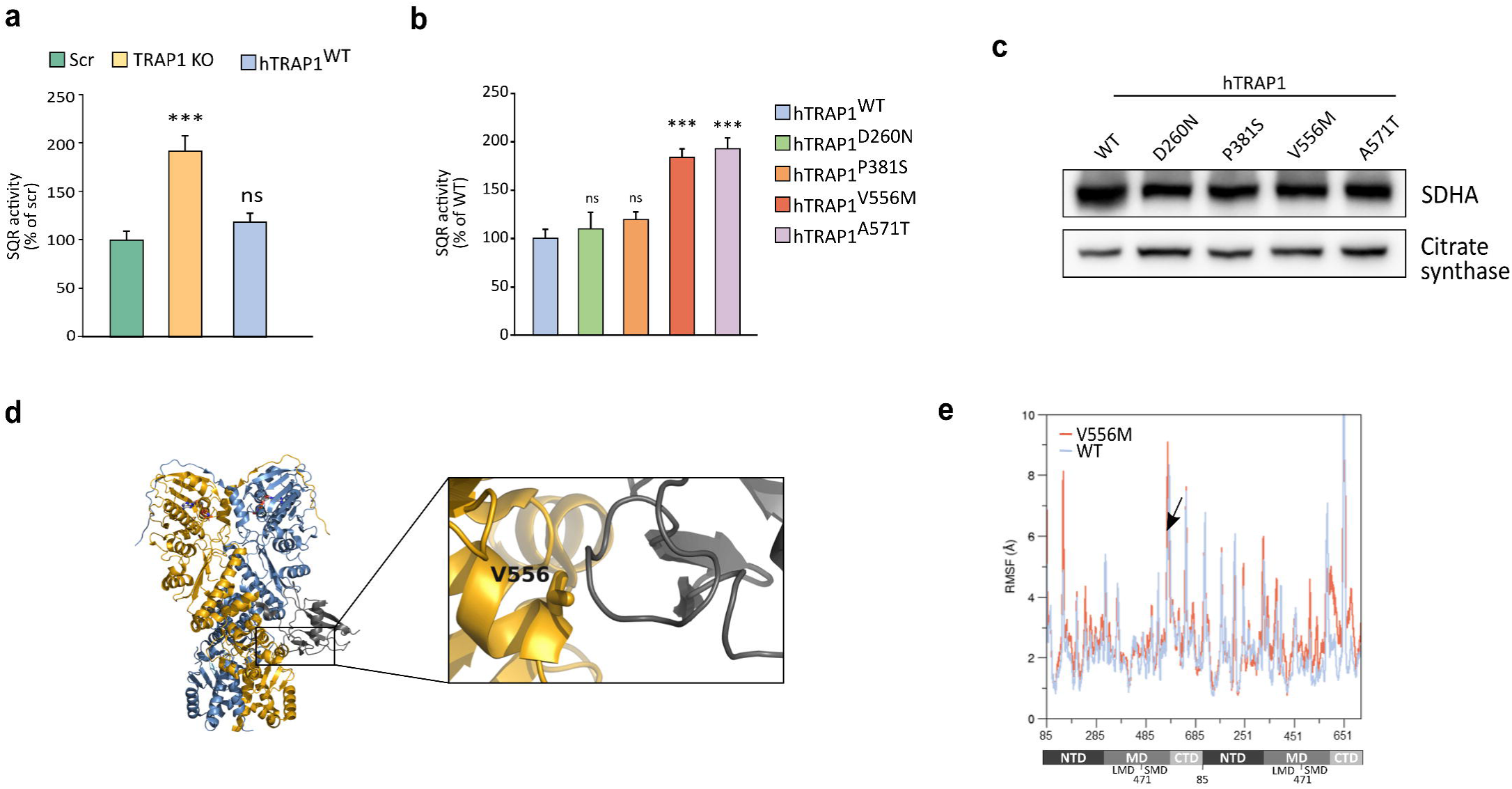
TRAP1 point mutations differentially affect succinate dehydrogenase activity. **a-b** Succinate dehydrogenase (SDH) activity measured in sMPNST scr, TRAP1 knockout cells, and in cells expressing either human wild-type or mutant forms of TRAP1. Data are reported as average ± S.E.M. of 3 independent experiments with a two-tail unpaired t test with each mutant compared to hTRAP1-WT expressing cells (**, p value < 0,01; ****, p value < 0,0001; n.s., not significant). **c** Western-blot of sMPNST cells expressing mutant form of TRAP1 showing basal expression of subunits A of Succinate Dehydrogenase. **d** Reproduction of PDB code 7KCM showing Trap1 bound to the model client protein SdhB (in black). Valine 556 is shown in stick and a zoomed view of its contact with SdhB is shown in the inset. **e** Root mean square fluctuation (RMSF) per residue considering the backbone atoms only. The mutated positions are indicated with an arrow. On the x axis numbering of zTrap1 is as in PDB code 4IPE and is shown together with the corresponding domains of prot. A and prot.B. For each protomer residue numbering is from 85 to 719.

Taken together our data suggest that two of the mutations under analysis impact TRAP1 chaperone activity, altering its ability to interact with and modulate SDH in a manner independent of its ATPase activity.

### Point mutations influence TRAP1 pro-neoplastic activity

We have previously found that TRAP1-mediated inhibition of SDH has a pro-neoplastic effect in various tumor cell models(16, 17). Hence, we tested the effect of TRAP1 variants in *in vitro* tumorigenic assays. Cultured sMPNST cells form foci overcoming contact inhibition, and this was strongly reduced upon genetic ablation or pharmacological TRAP1 inhibition with compound 5, and the tumorigenic potential of sMPNST cells was rescued upon re-expression of the h-TRAP1-WT (**Fig. 5a, Supplementary Fig. 5a-c and** (**17**)).

**Figure 5.**
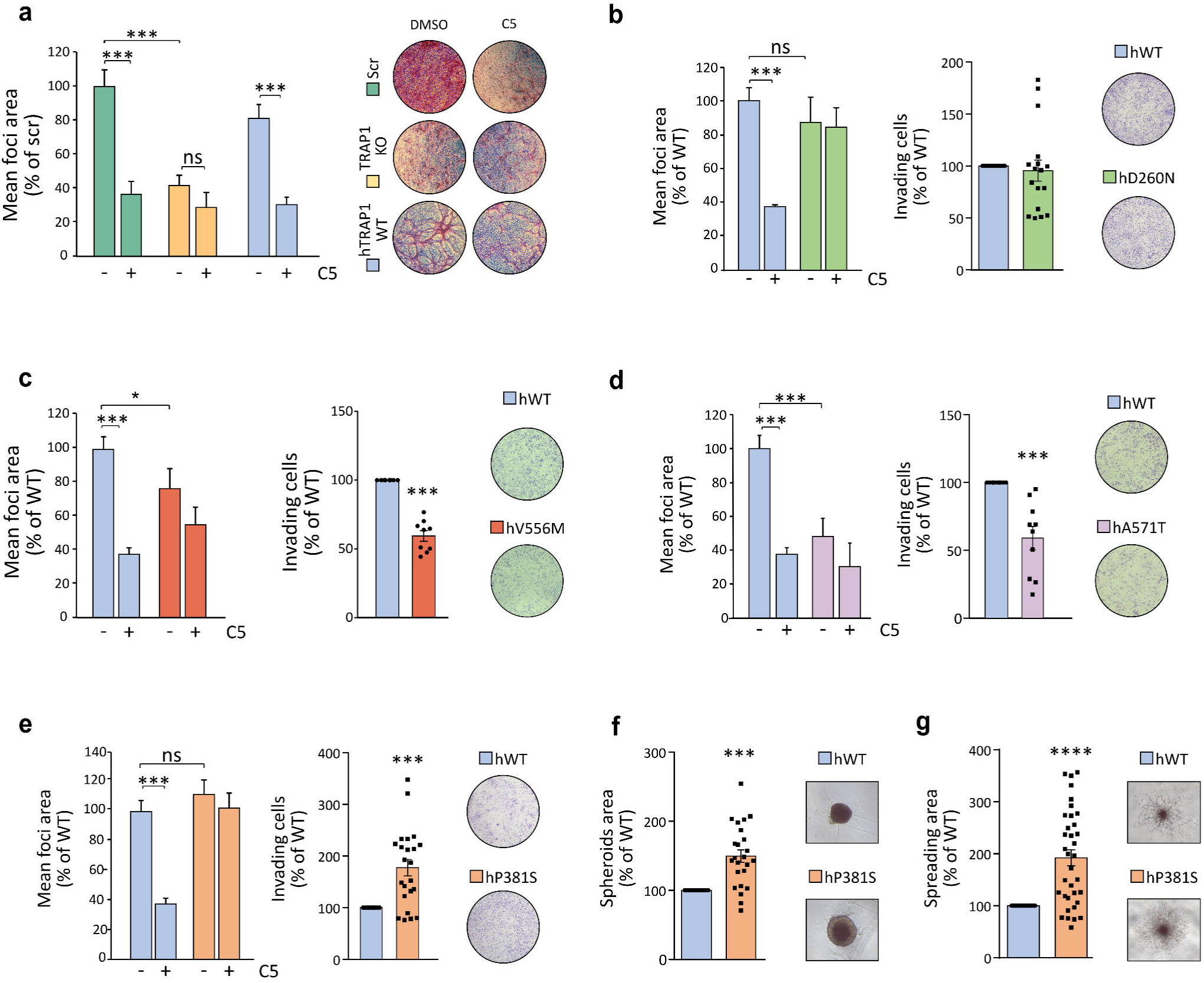
Effect of TRAP1 point mutations on its pro-neoplastic activity. **a** Focus-forming assay on sMPNST cells (scr, TRAP1 KO and re-expressing hWT-TRAP1) grown for 10 days with or without selective TRAP1 inhibitor compound 5 (25uM). Data are reported as mean of foci area normalized to scr SMPNST cells, and presented as mean ± SEM (n=3 independent experiments with 3 replicates for each one); ***p < 0.001 with one-way ANOVA with Bonferroni’s test. **b-e** Focus-forming and invasion assay on sMPNST cells expressing WT or TRAP1 mutants (**b** D260N, **c** V556M; **d** A571T; **e** P381S). Where indicated cells were treated with 25uM of compound 5. For focus forming assay, data are reported as mean of foci area normalized on WT-TRAP1 expressing cells. For invasion assay data are reported as area covered by invading cells normalized on WT-TRAP1 expressing cells. Data are presented as mean ± SEM of at least 3 independent experiments with 3 replicates for each one); ***p < 0.001 with one-way ANOVA with Bonferroni’s test for focus forming and t-test for invasion assay. **f** Spheroids formed by sMPNST expressing h-TRAP1-WT and h-TRAP1-P381S mutant. Spheroids area was measured after 10 days of sphere formation and quantified by ImagJ software. **g** Branching morphogenesis performed in sMPNST spheroids. Matrigel was added after 3 days of spheroids formation. The spreading of spheroids was estimated by measuring the area of branches using the imageJ software. Data are presented as mean ± SEM of at least 3 independent experiments with a t-test analysis, ***p < 0.001.

Notably, cells expressing the D260N mutant retain their ability to form foci and invade, while displaying resistance to compound 5 treatment (**Fig. 5b and Supplementary Fig. 5a,d**). In contrast, cells expressing the V556M and A571T mutants exhibit a diminished capacity for foci formation and invasion (**Fig. 5c-d and Supplementary Fig. 5 a, e and f**). Conversely, expression of hTRAP1-P381S markedly increases the *in vitro* tumorigenicity of MPNST cells, as they displayed greater capacity to form foci and to grow in 3D both as spheroids and in branching morphogenesis experiments (**Fig. 5e-g and Supplementary Fig. 5g**). It is worth noting that D260 and P381S mutations confer insensitivity to compound 5 treatments as cells continue to form foci to the same extent of control cells (**Fig. 5b, g and Supplementary Fig. 5d** and g).

These data highlight the biological relevance of TRAP1 point mutations, as they can tune its protumorigenic effects.

## Discussion

In this study, we provide insights into how specific point mutations affect TRAP1 structural stability, ATPase activity, bioenergetic and tumorigenic functions. These findings have crucial implications for the understanding of the structure-activity relationship of the chaperone and for advancing our comprehension of TRAP1’s role in disease pathogenesis. Additionally, understanding how structural changes impact protein functions could be essential for the development of highly selective TRAP1 inhibitors. The molecular chaperone TRAP1 has emerged as a multifaceted protein with a critical role in the maintenance of mitochondrial activities both in health and diseases. Some studies have linked TRAP1 point mutations to various pathological conditions, but how these mutations affect chaperone functions is unknown, particularly in the context of cancer, where TRAP1 has been extensively studied. Our high-throughput screening identified several point mutations in TRAP1 drawing from three protein databases: ClinVar, COSMIC and CBioPortal. The five variants analyzed in this study (D260N, P381S, V556M, A571T, and T600P) were selected according to both their position in protein sequence and MAVISp prediction regarding their effects on the structural stability and activity of the protein. Molecular Dynamics (MD) analyses show that these mutations significantly impact TRAP1’s conformational dynamics, with each substitution altering the local hydrogen bond network and affecting the protein’s internal dynamics, as measured by Distance Fluctuations (DFs) among residue pairs. DF assesses the dynamic coordination between residue pairs, with small CP values identifying mechanically connected amino acids that move cooperatively and significantly contribute to functional motions. The DF score to the mutated position (P_mutA/B_) changes significantly for cancer-related mutations, and the profile of internally connected residues along the sequence is altered compared to the one of the WT protein. Mutations such as D260N and P381S, associated with functional or hyper-functional TRAP1, increase the number of internally coordinated residues. In contrast, other mutations reduce internal coordination, making TRAP1 more flexible. Notably, P381S and V556M are in the M-domain near the client-binding region, suggesting that disrupted contacts with clients (as seen in the TRAP1-SDH complex) may directly impact client maturation and activation.

The variants D260N and A571T are predicted by the MAVISp modules to have neutral effects, suggesting that they could have neutral and/or passenger effects in cancer. D260 is distant from the client binding site and does not affect TRAP1 interaction with client proteins like SDH. Nonetheless, the alteration of the hydrogen bond network in the active site and the increased rigidity of the NTDs might explain the increase in ATPase activity and the ineffectiveness of compound 5. Our screening identified the P381S mutation as the one with the highest pathogenicity score. This mutation does not affect TRAP1 ATPase activity but significantly increases the tumorigenic potential of MPNST cells by enhancing foci formation and 3D growth, and it confers resistance to inhibitor compound 5. The V556M mutation is classified as neutral, with its predicted damaging effect limited to local alterations at the client binding site. Our results show increased flexibility around V556M associated with a significant reduction in the ability of TRAP1 to inhibit succinate dehydrogenase and in the tumorigenicity of cells expressing this mutant protein. A promising approach would be to test the effects of this variant on the recruitment of protein clients and whether the alteration of this region affects the TRAP1 chaperone activity. TRAP1 functions as a homodimer with the dimerization site in the C-terminal domain (CTD). The recurrent T600P variant within the CTD exhibits significant destabilizing effects, disrupting interactions with surrounding residues and the hydrogen bond network. This instability highlights the crucial role of CTD in maintaining TRAP1 structural integrity. Overall, our results point out the complex role of TRAP1 in cancer biology, showing that TRAP1 mutations have differential effects and revealing no direct correlation between its ATPase activity and functional effects. To exert its proteostatic functions, TRAP1 could form multimeric complexes with clients, whose composition and biological outputs could be flexibly adapted to the changes experienced by cells, with a particular importance in the highly unstable tumor microenvironment. Perturbing this central hub is expected to impair the formation of functional complexes with large-scale consequences that reverberate on mitochondrial and cellular homeostasis. Examining other variants identified in this study could enhance our understanding of TRAP1’s structure-activity relationships and aid in developing more selective and mutation-specific therapeutic strategies.

In conclusion, our study provides a comprehensive analysis of how specific point mutations affect TRAP1 structure, dynamics, function and role in tumorigenesis, underscoring its potential as a pharmacological target.

## Materials and Methods

### Analyses with the MAVISp computational framework

We applied the computational framework of MAVISp (Multi-layered Assessment of VarIants by Structure for proteins) to integrate pathogenicity prediction and molecular understanding of the effects of selected variants. We used the MAVISp framework (https://doi.org/10.1101/2022.10.22.513328) and referred to the RefSeq ID NP_057376 to retrieve information on the seven variants of interest, as well as other missense variants, reported in ClinVar (32) COSMIC (33, 34) and cBioPortal(35). The results can be retrieved from the MAVISp database (https://services.healthtech.dtu.dk/services/MAVISp-1.0/).

We applied the simple mode of MAVISp using the structure of the AlphaFold2 (AF2) model of human TRAP1 protomer retrieved on the 10th of October 2022 from the AF2 database. We verified the agreement between the model and known experimental structures and the overall structural quality of the model. We trimmed the model to remove regions with low pLDDT scores at the N-terminus (residues 1-93), including the region that is not present in any of the experimental structures available of TRAP1 (1–69) and the region that makes trans-protomer interactions (70–93). Although experimental structures of TRAP1 and the Alphafill database (36) contain data on the location of cofactors, including ATP, non-hydrolyzable analogs of ATP and magnesium (Mg^2+^) ions, we did not include any cofactor in the MAVISp assessment and calculations on the TRAP1 protomer, following the default protocol of MAVISp when applied for high-throughput purpouses. Thus, predictions regarding the proximity of cofactor binding sites should be made with caution. This includes the residues: E115, N119, A123, K126, D158, M163, N171, L172, R177-S180, G199-F205, and T251 evaluated looking at the surrounding from ANP and 4 Å cutoff of the structure of human TRAP1 fragment (PDB ID 5HPH)(37). Furthermore, we applied MAVISp to investigate the effects of variants on protein-protein interactions using a model of human TRAP1 homodimer (including residues 70-704) produced with a standalone version of Alphafold-Multimer(38). We verified the model quality by checking the PAE scores and the predicted interfaces with visual inspection with PyMOL. In the generated model, we included two Mg^2+^ ions, superimposed using their locations in the X-ray crystal structure of Danio rerio TRAP1 (PDB ID 4IPE)(28).

In addition to the standard modules provided by MAVISp [27], we also applied an additional module that allow to predict the local and long-range effects of variants on functional sites, which have been recently developed and applied for another case of study (39). In this analysis, we defined as “functional sites” the residues proposed to be part of the client binding site of TRAP1. We defined these residues by previous literature and analysis of i) the cryo-EM structure of human TRAP1 in complex with the client protein Succinate dehydrogenase B (PDB ID 7KCM)(40), ii) the X-ray crystal structure of zebrafish TRAP1 with the inhibitor Mitoquinone (PDB ID 7EXP,(41)) and iii) the binding sites of small molecules designed against an allosteric site in the middle domain of TRAP1(17).

In the STABILITY module of MAVISp, we estimated changes in folding free energy upon mutation using foldx5 with MutateX(42), Rosetta cartddg2020 (43) protocol and ref2015 (44) energy function with RosettaDDGPrediction (45) and the RaSP workflow(46). We used two consensus approaches among these methods to classify the variants, as explained in the original publication(42). For investigating the effects on protein-protein interactions in the TRAP1 homodimer we estimated changes in free energy calculations upon mutation using foldx5 with MutateX (42) and Rosetta flexddg protocol (43) and talaris2014 (44) energy function with RosettaDDGPrediction(45). We applied the LONG_RANGE module of MAVISp on the allosteric signaling map generated from AlloSigMA 2 (47) to predict changes in long-range structural communication and possible allosteric modulation. The details about the workflow applied are reported in the MAVISp publication(27). For the PTM module, we applied a decision tree to classify the effects of each variant implemented in the MAVISp framework(27). We collected the pathogenicity scores from DeMaSk(48), EVE(49), and AlphaMissense (50) included in MAVISp.

### Protein structures preparation and force field parametrization

The structure of WT zTrap1 was obtained from PDB code 4IPE(28), with missing loops modelled based on previous simulations(29, 51). Each zTrap1 protomer was reconstructed from Thr85 to His719. All Co^2+^ cations were deleted and the AMP-PNP ligand was edited to ATP by replacing the N_β_-H group with oxygen. Pymol 2.6 (Schrödinger, Inc) was employed to mutate the appropriate residues in both protomers to obtain the zTrap1 mutant models corresponding to the D260N, P381S, V556M, A571T and T600P hTrap1 mutants.

Subsequently, hydrogens were introduced using the utility Reduce (AmberTools21)(52), while protonation/tautomerization states were assigned using propka3 (53), with Histidine residues modelled as described in previous reports(51). For side chains with alternate locations we retained the most abundant conformer; accordingly, Cys516 and Cys542 were kept in the reduced oxidation state with no disulfide bridge. Also, the N- and C-termini were capped with acetyl (ACE) and N-methyl (NME) fragments respectively. Protein residues were parametrized using the Amber forcefield ff99SB (54). For ATP we adopted the parameters published Meagher and coworkers(55), whereas for Mg^2+^ those reported by Allnér et al.(56). Tleap (AmberTools21) (52) was employed for building a truncated octahedral box of water (TIP3P model) (57) with a distance of at least 10 Å between every protein atom and the closest edge. Sodium cations were added to neutralize the charge and were modelled using parameters by Joung and Cheatham(58).

### MD preproduction stages

Atomistic molecular dynamics simulations were conducted using the sander and pmemd modules from AmberTools (version 21) and Amber (version 20) (52), respectively. The sander engine was utilized for the minimization, solvent equilibration, and heating stages. The GPU-accelerated pmemd.cuda engine was employed for the later stages of equilibration and for the production(59). For all systems, three independent replicas were run comprising two minimization steps, heating, system equilibration and production. In the first minimization round (500 steps of steepest descent + 500 steps of conjugate gradient) only the solvent was unrestrained while all other residues (protein residues, ATP and Mg^2+^) were subjected to a harmonic positional restraint with a 500 kcal mol^-1^ Å^−2^ constant. Afterwards, the whole system was allowed to relax through 1000 steps of steepest descent and 1500 of conjugate gradient minimization. The cut-off for the calculation of Lennard-Jones and Coulomb interactions was set to 8 Å, beyond, only Coulomb interactions are computed in direct space up to 10 Å, shifting thereafter to the particle mesh Ewald method(60).

Minimization is followed by random assignment of initial velocities to match a temperature of 25 K, and a rapid heating to 300 K (20 ps; NVT ensemble) in which positions of non-solvent molecules are gently restrained with a 5 kcal mol^−1^ Å^−2^ constant. The Langevin thermostat (61) is employed for temperature control with a weak coupling at this stage (0.75 ps^−1^ collision frequency). SHAKE constraints for bonds containing hydrogens were introduced and kept thereafter(62). Accordingly, the time step was set to 2 fs and will remain so for the rest of the simulation. Subsequently, all restraints were released, and the system was equilibrated at 300K for 1 ns in the NpT ensemble to allow for density adjustment. Pressure control was enforced through the Berendsen’s barostat (63) with a 2 ps relaxation time while coupling to the Langevin thermostat was increased to 1 ps^-1^ collision frequency.

### MD production in NVT ensemble and meta-trajectory of the equilibrated portions

All systems were simulated for 300 ns in NVT ensemble during the production run of each replica, with coupling to the thermostat tightened (5 ps^-1^ collision frequency). Coordinates were printed every 10 ps for a total of 30000 frames for each replica trajectory. To consider only equilibrated portions of each replica, we kept only the last 200 ns of each trajectory with one every 5 frames retained for further analysis, resulting in a total of 4000 frames per replica with a spacing of 50 ps. For each Trap1 protein system, MD trajectories from the three replicas were concatenated to form meta-trajectories using cpptraj (AmberTools 21)(52); in doing so, we centred and imaged the trajectory, stripped solvent and Na^+^ ions, and aligned all structures to the first frame in the first replica using, as reference, backbone heavy atoms of residues involved in secondary structures. As a result, meta-trajectories of 12000 frames covering a time interval of 600 ns were obtained for each Trap1 system.

### Trajectory analysis: H-bond interactions, Secondary Structures and Root Mean Square Fluctuation (RMSF)

All the analyses were carried out on the entire 600 ns meta-trajectory employing *cpptraj* (AmberTools, version 21) (52). The nature and occupancy of H-bond interactions involving the mutated positions were analysed using the h-bond command leaving default settings for H-bond recognition (angle cutoff = 135°, distance cutoff = 3.0 Å). Calculation of secondary structure content for residues of interest was carried out with the command *secstruct*. Moreover, to evaluate relevant variations in the flexibility of residues upon mutation we computed the per-residue Root Mean Square Fluctuation (RMSF) considering only backbone atoms and taking the average structure of the meta-trajectory as reference.

### Distance fluctuations analysis (DF)

Distance fluctuation analysis was carried out to investigate dynamic allosteric networks. This method was developed internally and its significance for describing the internal dynamics of a protein was extensively reported (7, 64, 65). A brief overview is provided here. The DF matrix was calculated considering only Cα along the entire 600 ns long meta-trajectory for each zTrap1 protein system. This is a N×N matrix (N= number of residues) in which every element corresponds to a pairwise DF parameter (score) given by the following equation:

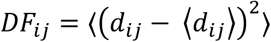

Here, d*_ij_* is the time-dependent distance between the Cα of *i* and *j* amino acids and the pointy brackets indicate averaging over the MD meta-trajectory. The DF score, in practice a variance of residue pairwise distances, computed for the Cα of every residue pair, quantifies the degree of mechanical coordination between the examined aminoacids. In particular, small DF scores can be associated with regions characterized by a high degree of mechanical coordination and moving quasi-rigidly, whereas high scores identify flexible regions moving in an uncoordinated fashion.

Going one step further, one can identify for each residue in the sequence the identity and quantity of the residues that are highly coordinated to it. To do so we set as a threshold for the DF score, the average value of the Local Fluctuation for each protein (LF). The Local Fluctuation for a residue i can be calculated by averaging the DF_ij_ values between *i* and every neighbouring j residue with *j=i±2* along the sequence. At this point, residues at an average distance higher than 5Å and with a DF score lower than LF can be considered to be mechanically connected. The number of residues connected to each position, termed mechanical connectivity, will be indicated with η and the variation between η_mut_ and η_WT_ is an indication of changes in allosteric communication upon mutation (Δη_mut_).

Another way for simplifying the evaluation of the impact of the mutation on the DF parameters is computing the DF difference matrix ΔDF_mut_ = DF_mut_ – DF_WT_ and normalizing it to obtain percentage differences taking DF_WT_ as a reference, thus obtaining a percentage difference matrix %ΔDF_mut_. In this matrix, negative values indicate residues that acquire mechanical coordination (rigidity) upon mutation (lower DF scores in the mutant); whereas positive values identify residues that lose mechanical coordination upon mutations (higher DF scores in the mutant). It is also possible to extract the columns relative to mutated positions of prot. A or B and project it onto the protein 3D structures. These projections, which represent the variations in the DF scores of all residues to the mutated position, will be termed P_mutA_ or P_mutB_ depending on the protomer from which the projected column of the %ΔDF matrix was extracted.

### Analysis of TRAP1 expression

GeneCards, GTEx Portal and Gene Expression Atlas were used to retrieve TRAP1 expression data at cell and tissue level. The GENT2 database was used to collect expression data from cancer samples. Somatic TRAP1 variants found in cancer were retrieved from COSMIC and mapped on the TRAP1 3D-structure using Chimera. Prediction of residue-residue interactions lost upon mutations was performed with RING2.0.

### Cell maintenance

Malignant peripheral nerve sheath tumor cells (sMPNST cells) were established from neurofibromin 1 (Nf1)-deficient skin precursors (SKP) and were kindly provided by Dr. Lu Q. Le, University of Texas Southwestern Medical Center, Dallas, TX. sMPNST and HEK 923 cells were grown Dulbecco’s Modified Eagle’s Medium (DMEM) and RPMI-1640 supplemented with 10% fetal bovine serum (FBS) 100 units/ml and 1% penicillin and streptomycin. All cells were cultured at 37°C in a humidified atmosphere containing 5% CO_2_.

### Generation of mutant cells

The QuickChange site-directed mutagenesis kit was used to generate mutant forms of human TRAP1 (hTRAP1-WT; hTRAP1-D260N; hTRAP1-P381S; hTRAP1-V556M; hTRAP1-A571T; hTRAP1-T600P). pBABE vectors (Addgene Plasmid #1767) containing both Wild-Type or mutant variants form of hTRAP1 were used to stably transfect TRAP1 knock-out sMPNST cells previously generated by Sanchez et al. pBABE vectors were co-transfected with packaging plasmids PMD2.G and psPAX2 into HEK 293 cells for viral production. Recombinant virus was used to infect sMPNST which were subsequently selected with 0.75 mg/ml G418 (sigma).

### Production of recombinant proteins

The Wild-type or mutant forms of mature sequence of human TRAP1 without the mitochondrial targeting sequence were cloned into the pRSET-Sumo plasmid to obtain a N-terminally 6xHis-Sumo-tagged TRAP1 fusion protein. Recombinant TRAP1 (WT and mutants) was produced in BL21(DE3) E. coli cells after 18h induction with 0.4 mM IPTG. After bacteria lysis, the soluble fraction was incubated with Ni-NTA resin (Merk # GE17-5268-001) according to manufacturer indications. Proteins release from the resin packed in a FPLC column was achieved using a buffer composed by 50 mM potassium phosphate at pH 7.0, 300 mM sodium chloride, 500 mM imidazole and 3 mM b-mercaptoethanol. The eluted protein was dialyzed overnight at 4° C in a dialysis buffer composed of 10 mM Tris-HCl pH 8.0, 200 mM NaCl, 1 mM b-mercaptoethanol and then filtered and maintained at 4°C.

### TRAP1 ATPase activity

TRAP1 ATPase activity was measured by quantifying the release of inorganic phosphate as PO4 nmoles/TRAP1 nmoles/min. The ATPase assay was performed at 37 °C for 1 h, incubating 5 μg of proteins with 200 μM ATP and 10 mM MgCl2, in the reaction buffer, in the dark. The amount of inorganic phosphate (PO_4_^3-^) released due to the activity of TRAP1 was measured using the Malachite Green Phosphate Assay Kit (AMBION), by detecting the increase in absorbance at 650 nm with a plate reader.

### Protein isolation and western blot analysis

For Western immunoblot analyses, cells were lysed at 4 °C in an RIPA buffer (Tris-HCl 50 mM pH7.4, NaCl 150 mM, NP40 1%, sodium deoxycholate 0.5%, SDS 0.1%, EDTA 2 mM and protease inhibitors (Sigma)) and then clarified at 14,000 rpm for 30 min at 4 °C. Protein quantification was carried out by using a BCA Protein Assay Kit (Thermo-Scientific). Extracted proteins were boiled at 50 °C with Laemli buffer for 5 min, separated in reducing conditions by using NuPage Novex 4%-12% Bis-Tris gels (Life Technologies) and transferred in Hybond-C Extra membranes (Amersham). Primary antibodies were incubated for 16 h at 4 °C (TRAP1 Santa Cruz #sc-73604, Citrate Synthase Abcam #ab96600). Proteins were visualized using the UVITEC imaging system following incubation with horseradish peroxidase-conjugated secondary antibodies.

### Measurement of succinate:coenzyme Q reductase (SQR) activity of SDH

Succinate dehydrogenase (SDH) activity was measured as already described by Sanchez et al. Briefly, samples were collected and lysed at 4°C in a buffer composed of 25 mM potassium phosphate, pH 7.2, and 5 mM magnesium chloride containing protease and phosphatase inhibitors. Total lysate was quantified with BCA protein assay Kit (Thermo-Scientific), and 40 ug of protein per trace was incubated 10 min at 30 °C in the presence of 20 mM sodium succinate and 10 mM alamethicin. After incubation, a mix composed of sodium azide 5 mM, Antimycin A 5 μM, Rotenone 2 μM, and Coenzyme Q1 65 μM was added. SDH activity was measured by following the reduction of 2–6 dichlorophenolindophenol (DCPIP) at 600 nm (ε = 19.1 nM−1 cm−1) at 30°C, each measurement was normalized for protein amount.

### *In vitro* tumorigenic assay

Focus forming assays were performed on sMPNST cells grown in 12-well culture plates in DMEM medium supplemented with 10% fetal bovine serum. When cells reached sub-confluence, serum concentration was decreased to 1%. After 10 days, plates were washed in PBS, fixed in methanol for 30 min, stained with GIEMSA solution for 1h and analyzed with ImageJ software.

### Spheroids formation and *in vitro* migration assay

For spheroid generation, 5000 sMPNST cells were plated in round-bottom 96 well plates, centrifuged at 100g for 1 min at room temperature and incubated at 37°C 5% CO2. After 72h, 50ul of DMEM was replaced with 50ul of Matrigel, centrifuged at 100g a for 1 minute at 4°C and incubated at 37°C 5% CO2. After 5 days cell spreading was analyzed with ImagJ software.

### Data availability

The aggregated csv data from MAVISp are available at the MAVISp database: https://services.healthtech.dtu.dk/services/MAVISp-1.0/. Other data from MAVISp are deposited in the OSF repository. An overview of TRAP1 and its results from MAVISp is reported at: https://elelab.gitbook.io/mavisp/proteins/trap1.

### Statistical analysis

Data were analyzed and presented as mean ± standard deviation or standard error of the mean (SEM) as indicated in figures. Pairs of data groups were analyzed using paired and unpaired two-tailed Student’s *t* tests. In the case of more than two groups, a one-way analysis of variance (ANOVA) followed by Bonferroni post hoc test was applied. Statistical significance was determined using GraphPad Prism 8. Outliers were removed from analysis and were calculated by applying the outlier formula. Results with a *p* value lower than 0.05 compared to controls were considered significant and indicated as ^∗∗∗^*p* < 0.001, ^∗∗^*p* < 0.01, ^∗^*p* < 0.05. Each experiment was repeated at least three times.

## Acknowledgments

We thank Elena Trevisan and Marco Ardina for excellent technical assistance.

## Author contribution

C.L.: conceptualization, visualization, methodology, investigation, formal analysis, writing-original draft, writing-review and editing, supervision; A.M, M.B.: investigation, methodology, formal analysis; F.G, M.L.: methodology, formal analysis, writing-original draft; G.F, L.F, M.L.S, E.M.: methodology, formal analysis; G.C.: conceptualization, methodology, formal analysis; E.P.: conceptualization, methodology, formal analysis; A.R.: conceptualization, writing-original draft, writing-review and editing, funding acquisition, supervision.

---

**Supplementary Fig 1.**
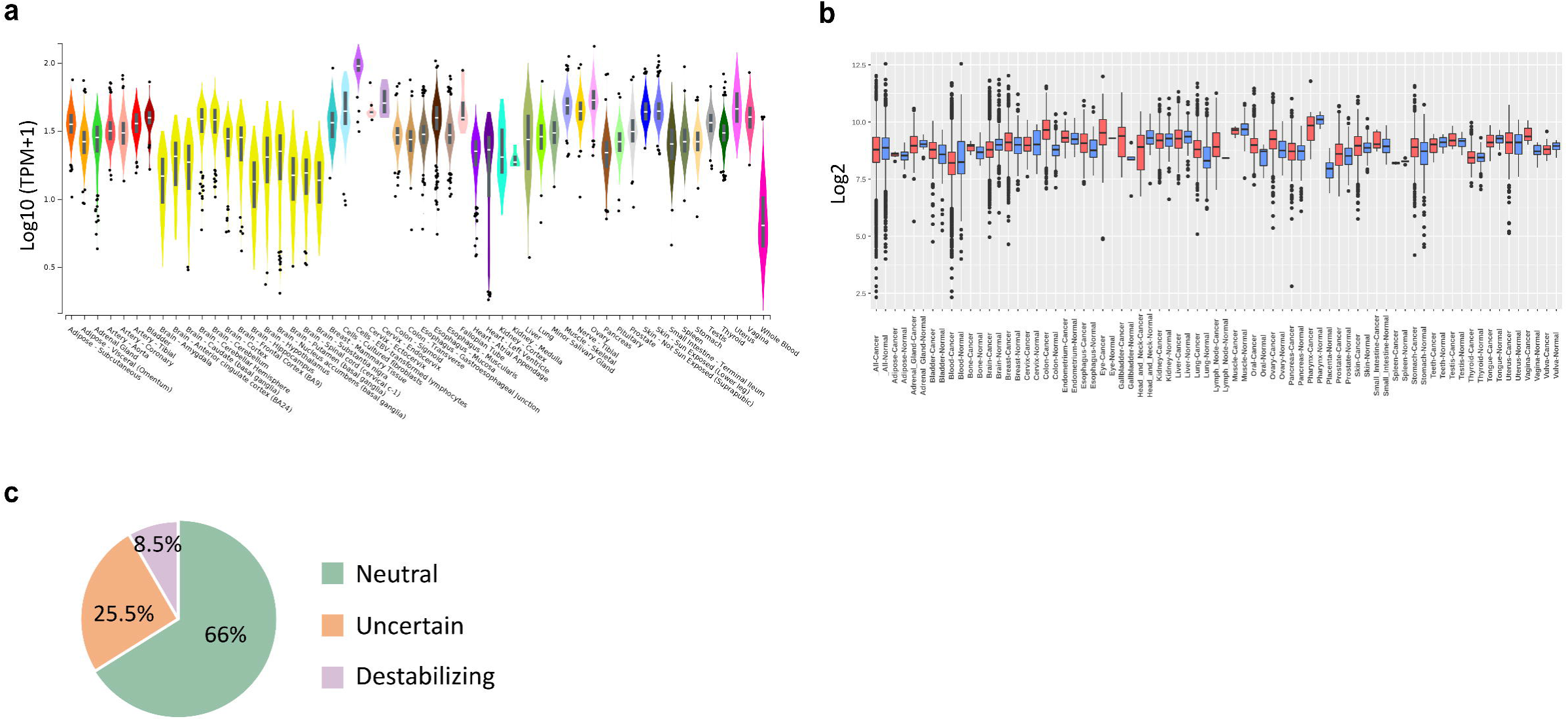
High throughput screening and classification of TRAP1 point mutations. **a** Box plot of human TRAP1 (ENSG00000126602.10) gene expression data from GTEx Portal. Expression values are expressed as logarithm of transcripts per million (TPM). Boxplots are expressed as median and 25^th^ and 75^th^ percentiles. Dots are for outliers (1.5 times above or below the interquartile range). **b** Comparison between TRAP1 expression in cancer and healthy samples. Red boxplots are for cancer sample, while blue boxplots are for healthy tissue. Dots are for outliers (1.5 times above or below the interquartile range). Expression data calculated using the Affymetrix HG-U133Plus2 dataset (GPL570) as reported in the GENT2 database. **c** classification of 310 TRAP1 variants retrieved on MAVISPs according to their effect on protein stability in neutral (66%), uncertain (25,5%) and destabilizing (8,5).

**Supplementary Figure 2.**
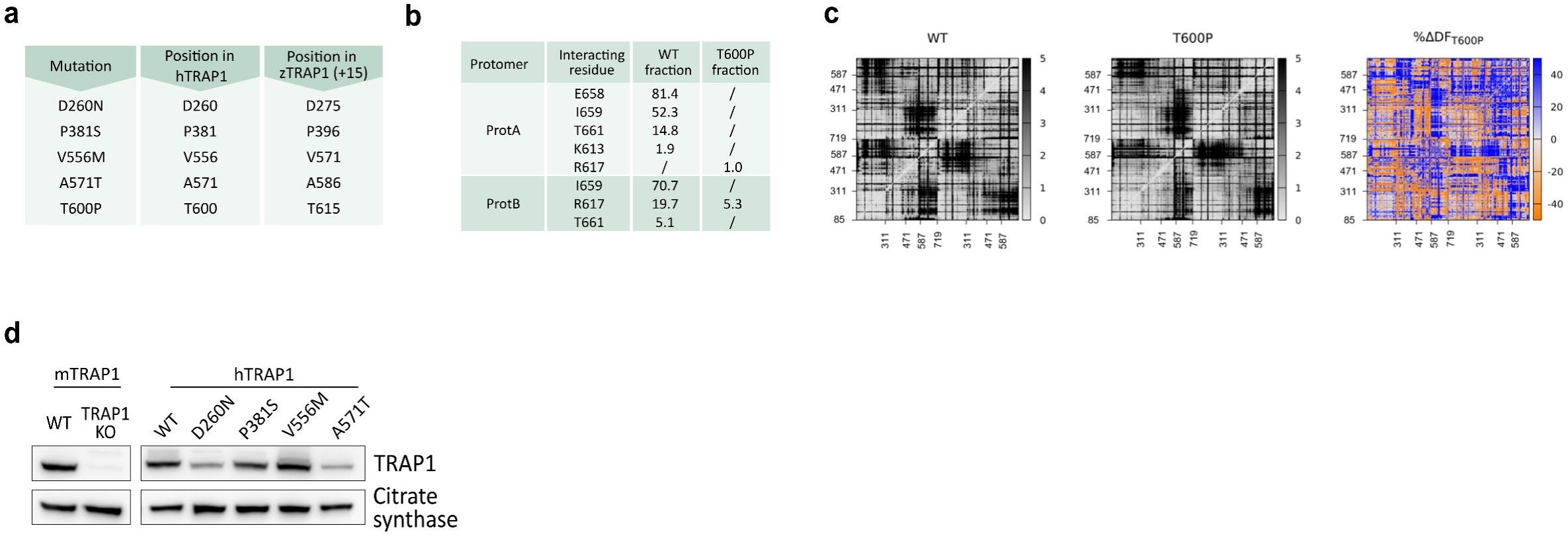
Effect of point mutations on TRAP1 stability. **a** Table indicating the position of each mutation in both human and Zebrafish TRAP1. In Zebrafish TRAP1 protein residue numbering is offset by 15 residues relative to the human protein. **b** Occupancy of H-bond interactions involving residue 615 (600 in hTrap1) in prot. A or B in WT and T600P zTrap1 during MD trajectories (12000 frames, 600ns). Numbering of residues is that of zTrap1 sequence as in PDB code 4IPE. To convert to hTrap1 numbering 15 should be subtracted. **c** Pairwise Cα Distance Fluctuation (DF) matrices of WT and mutant T600P-TRAP1 together with the percentage difference matrix %ΔDF. In DF matrices DF scores are coloured according to a grey scale as indicated in the colour bars. In the %ΔDF_mut_ blue areas (positive values) correspond to lower mechanical coordination in Trap1 mutant than in the WT protein whereas orange ones (negative values) indicate a higher coordination in the first. Grey/white areas are those unaffected by the mutation. On the x axis numbering is as in PDB code 4IPE and the values indicated correspond to the domain division. For each protomer residue numbering is from 85 to 719 and the assignment of domains is as follows: N-terminal Domain (NTD) residues 85-310; Middle Domain (MD), divided in the subdomains Large Middle Domain (LMD), residues 311-470 and Small Middle Domain (SMD) residues 471-586; C-Terminal Domain (CTD) residues 587-719. **d** Western-blot of scramble and TRAP1 KO sMPNST (left) re-expressing human wild-type and mutant forms of TRAP1 (right).

**Supplementary Figure 3.**
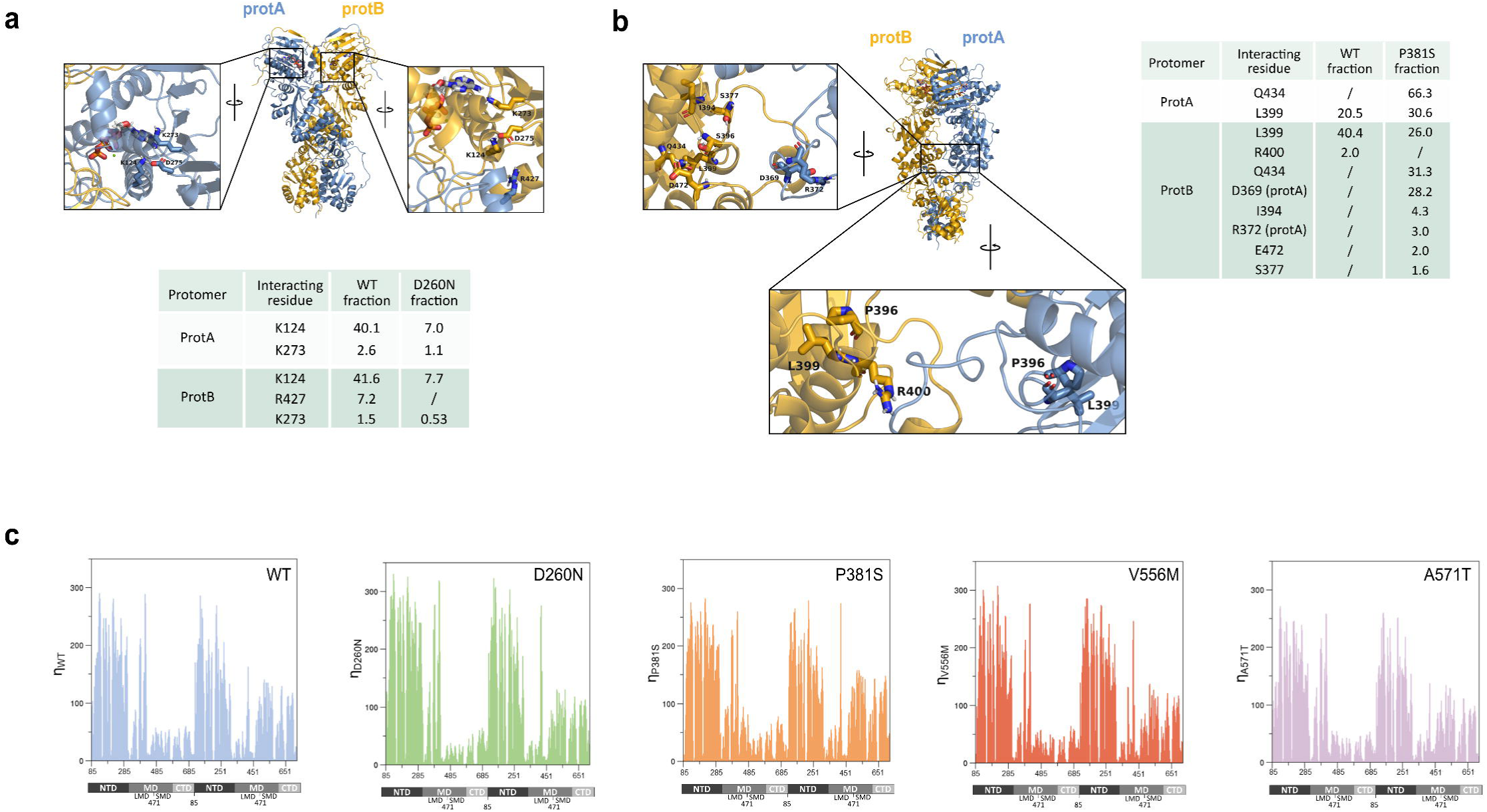
Effect of point mutations on TRAP1 molecular dynamics. **a** Occupancy of H-bond interactions involving residue 275 (260 in hTrap1) in prot. A (light blue) or B (dark yellow) in WT and D260N zTrap1 during MD trajectories (12000 frames, 600ns). Numbering of residues is that of zTrap1 sequence as in PDB code 4IPE. To convert to hTrap1 numbering 15 should be subtracted. **b** Structure of WT zTrap1 with P396 (P381 for hTrap1) and interacting residues listed in table shown as sticks together with ATP. Numbering is relative to zTrap1 sequence as in PDB code 4IPE. To convert to hTrap1 numbering 15 should be subtracted. **c** Mechanical connectivity index for each mutant along the sequence (η_mut_). On the x axis numbering of zTrap1 as in PDB code 4IPE is shown together with the corresponding domains of prot. A and B. For each protomer residue numbering is from 85 to 719 and the division in domains is as follows: N-terminal Domain (NTD) residues 85-310; Middle Domain (MD), divided in the subdomains Large Middle Domain (LMD), residues 311-470 and Small Middle Domain (SMD) residues 471-586; C-Terminal Domain (CTD) residues 587-719.

**Supplementary Figure 4.**
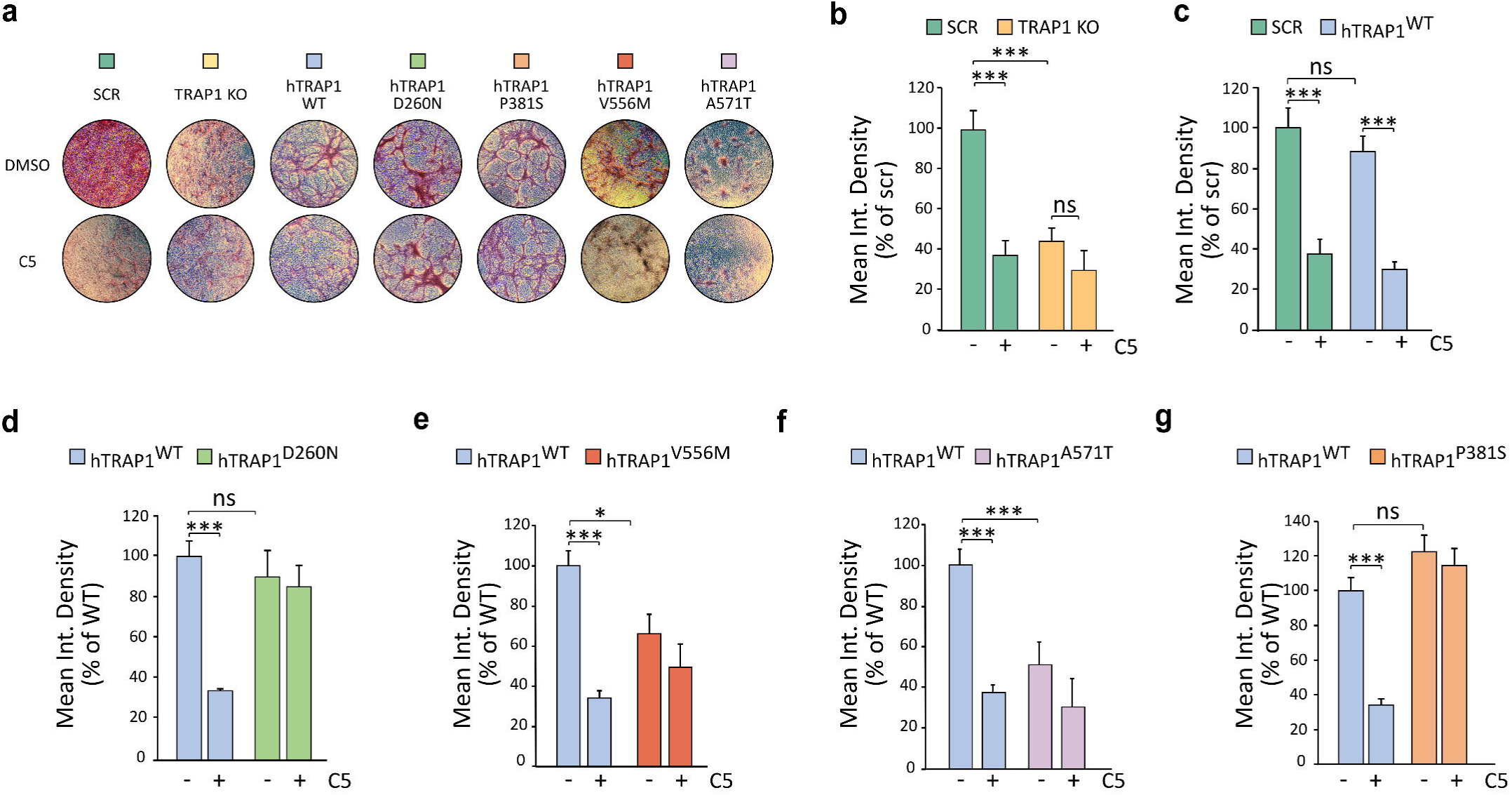
Effect of TRAP1 point mutations on its pro-neoplastic activity. **a** representative pictures of foci formed by sMPNST cells. **b-g** Focus-forming assay in SCR and TRAP1 KO sMPNST cells (**b-c**) or expressing h-TRAP1-WT or mutant forms (**d-g**). Cells were grown for 10 days with or without selective TRAP1 inhibitor compound 5 (25uM). Foci are quantified using an integrated density parameter that evaluates both their surface and thickness. Data are presented as mean ± SEM of at least 3 independent experiments with 3 replicates for each one; ***p < 0.001 with one-way ANOVA with Bonferroni’s test.

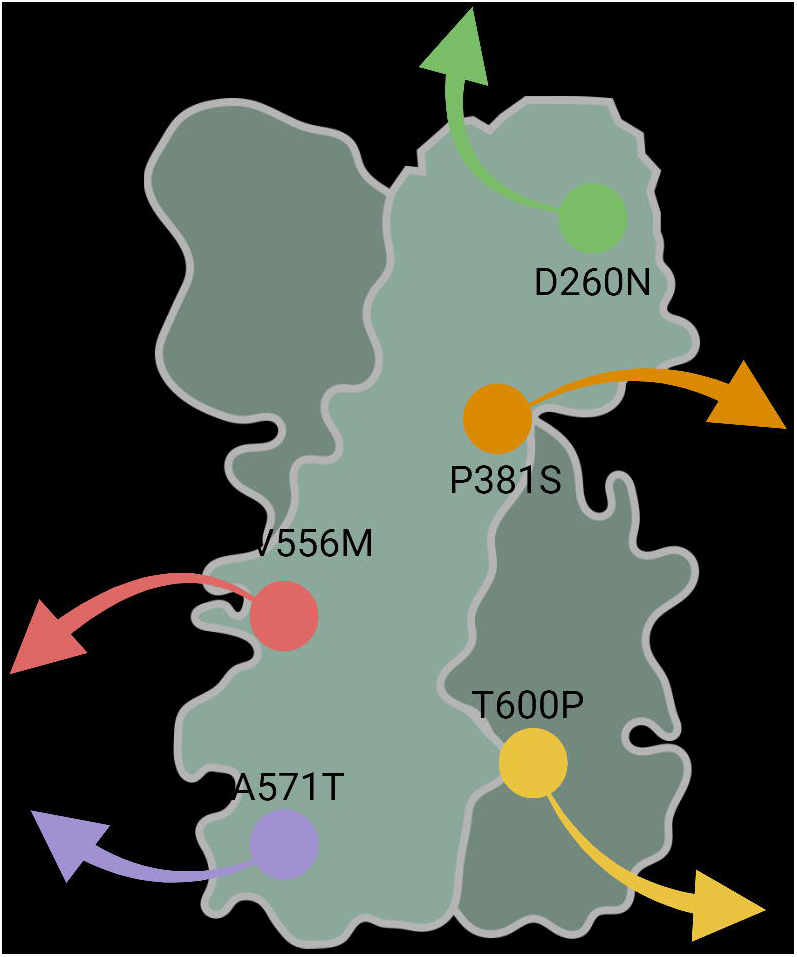

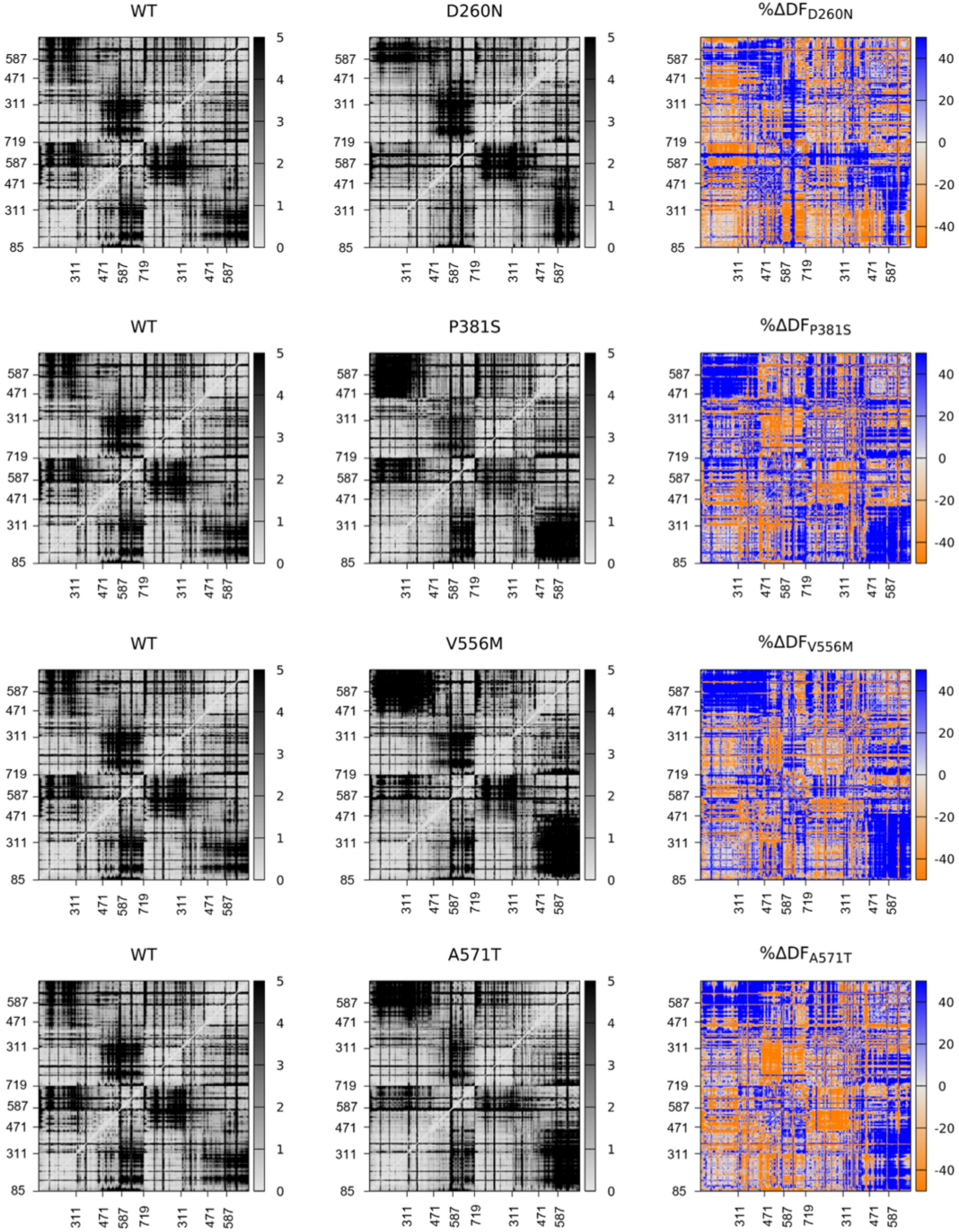

